# Preliminary estimation of the basic reproduction number of novel coronavirus (2019-nCoV) in China, from 2019 to 2020: A data-driven analysis in the early phase of the outbreak

**DOI:** 10.1101/2020.01.23.916395

**Authors:** Shi Zhao, Qianyin Lin, Jinjun Ran, Salihu S Musa, Guangpu Yang, Weiming Wang, Yijun Lou, Daozhou Gao, Lin Yang, Daihai He, Maggie H Wang

**Affiliations:** JC School of Public Health and Primary Care, Chinese University of Hong Kong, Hong Kong, China; Shenzhen Research Institute of Chinese University of Hong Kong, Shenzhen, China; Michigan Institute for Data Science, University of Michigan, Ann Arbor, Michigan, USA; School of Public Health, Li Ka Shing Faculty of Medicine, University of Hong Kong, Hong Kong, China; Department of Applied Mathematics, Hong Kong Polytechnic University, Hong Kong, China; Department of Orthopaedics and Traumatology, Chinese University of Hong Kong, Hong Kong, China; SH Ho Scoliosis Research Lab, Joint Scoliosis Research Center of Chinese University of Hong Kong and Nanjing University, Hong Kong, China; School of Mathematics and Statistics, Huaiyin Normal University, Huaian, China; Department of Mathematics, Shanghai Normal University, Shanghai, China; School of Nursing, Hong Kong Polytechnic University, Hong Kong, China

## Abstract

**Backgrounds:** An ongoing outbreak of a novel coronavirus (2019-nCoV) pneumonia hit a major city of China, Wuhan, December 2019 and subsequently reached other provinces/regions of China and countries. We present estimates of the basic reproduction number, *R*_0_, of 2019-nCoV in the early phase of the outbreak.

**Methods:** Accounting for the impact of the variations in disease reporting rate, we modelled the epidemic curve of 2019-nCoV cases time series, in mainland China from January 10 to January 24, 2020, through the exponential growth. With the estimated intrinsic growth rate (*γ*), we estimated *R*_0_ by using the serial intervals (SI) of two other well-known coronavirus diseases, MERS and SARS, as approximations for the true unknown SI.

**Findings:** The early outbreak data largely follows the exponential growth. We estimated that the mean *R*_0_ ranges from 2.24 (95%CI: 1.96-2.55) to 3.58 (95%CI: 2.89-4.39) associated with 8-fold to 2-fold increase in the reporting rate. We demonstrated that changes in reporting rate substantially affect estimates of *R*_0_.

**Conclusion:** The mean estimate of *R*_0_ for the 2019-nCoV ranges from 2.24 to 3.58, and significantly larger than 1. Our findings indicate the potential of 2019-nCoV to cause outbreaks.

## Introduction

The atypical pneumonia case, caused by a novel coronavirus (2019-nCoV), was first reported and confirmed in Wuhan, China in December 31, 2019 [1]. As of January 26 (17:00 GMT), 2020, there have been 2033 confirmed cases of 2019-nCoV infections in mainland China, including 56 deaths [2]. The 2019-nCoV cases were also reported in Thailand, Japan, Republic of Korea, Hong Kong, Taiwan and the US, and all these cases were exported from Wuhan, see WHO news release https://www.who.int/csr/don/en/ from January 14-21. The outbreak is still on-going. A recently published preprint by Imai *et al.* estimated that a total of 1723 (95%CI: 427-4471) cases of 2019-nCoV infections in Wuhan had onset of symptoms by January 12, 2020 [3]. The likelihood of travel related risks of disease spreading is suggested by [4], which indicates the potentials of regional and global spread [5].

To the best of our knowledge, there is no existing peer-reviewed literature quantifying the transmissibility of 2019-nCoV as of January 22, 2020. In this study, we estimated the transmissibility of 2019-nCoV via the basic reproduction number, *R*_0_, based on the limited data in the early phase of the outbreak.

## Methods

We obtained the number of 2019-nCoV cases time series data in mainland China released by Wuhan Municipal Health Commission, China and National Health Commission of China from January 10 to January 24, 2020 from [6]. All cases were laboratory confirmed following the case definition by national health commission of China [7]. Although the date of submission of this study is January 26, we choose to use data up to January 24. Please note that the data of the most recent few days contain a number of infections that were infected outside Wuhan due to travel, and thus this part of infections is excluded from the analysis.

Although there were cases confirmed on or before January 16, the official diagnostic protocol was released by WHO on January 17 [8]. To adjust the impact of this event, we considered a time-varying reporting rate that follows a linear increasing trend, motivated by the previous study [9]. We assumed that the reporting rate, r(t), started increasing since January 17, and stopped at the maximal level at January 21. The time window of the reporting rate increase accounts for the official report on improving the 2019-nCoV surveillance released by the Chinese government by the end of January 21 [10], which also coincides with the average of the incubation periods of two other well-known coronavirus diseases, i.e., the middle east respiratory syndrome (MERS) and the severe acute respiratory syndrome (SARS), i.e., 5 days [11–13]. Denoting the daily reported number of new cases by *c*(*t*) for the *t*-th day, then the adjusted cumulative number of cases, 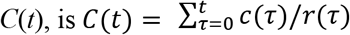. Instead of finding the exact value of *r*(*t*), we calculated the fold change in *r*(*t*) that is defined by the ratio of *r* on January 10 over that on January 24 minus 1. We illustrated six scenarios with 0- (no change), 0.5-, 1-, 2-, 4- and 8-fold increase in reporting rate, see Fig 1(a), (c), (e), (g), (i) and (k).

**Figure 1.**
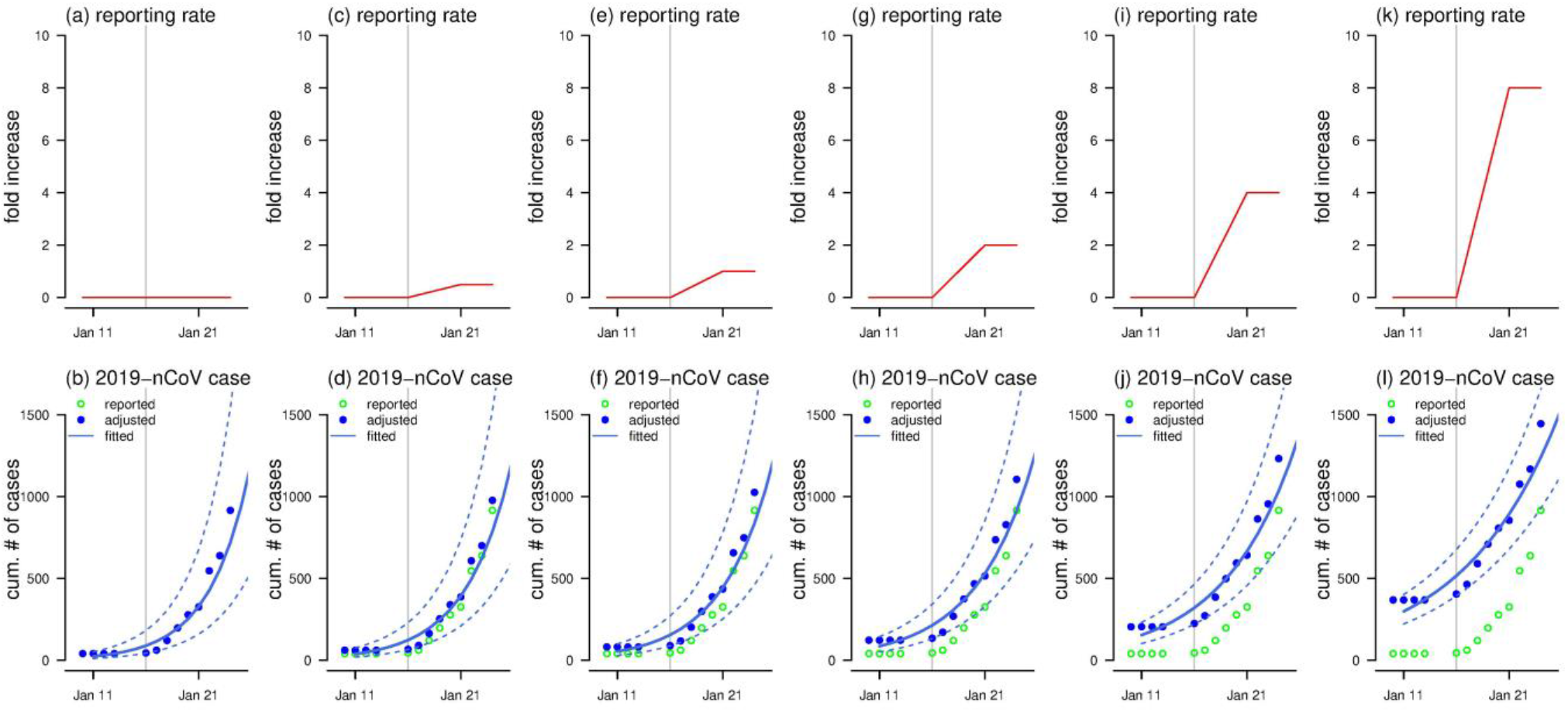
The scenarios of the change in the reporting rate (top panels) and the exponential growth fitting (bottom panels). The top panels, i.e., (a), (c), (e), (g), (i) and (k), show the assumed change in the reporting rate. The bottom panels, i.e., (b), (d), (f), (h), (j) and (l), show the reported (or observed, green circles), adjusted (blue dots) and fitted (blue curve) number of 2019-nCoV infections, and the blue dashed lines are the 95%CI. The vertical grey line represents the date of January 16, 2020, after which the official diagnostic protocol was released by WHO [8]. Panels (a) and (b) show the scenarios that the reporting rate was unchanged. Panels (c) and (d) show the scenarios that the reporting rate increased by 0.5-fold. Panels (e) and (f) show the scenarios that the reporting rate increased by 1-fold. Panels (g) and (h) show the scenarios that the reporting rate increased by 2-fold. Panels (i) and (j) show the scenarios that the reporting rate increased by 4-fold. Panels (k) and (l) show the scenarios that the reporting rate increased by 8-fold.

Following previous studies [14, 15], we modelled the epidemic curve obeying the exponential growth. The nonlinear least square (NLS) framework is adopted for data fitting and parameter estimation. The intrinsic growth rate (*γ*) of the exponential growth was estimated, and the basic reproduction number could be obtained by *R*_0_ = 1/*M*(-*γ*) with 100% susceptibility for 2019-nCoV at this early stage. The function *M*(·) is the Laplace transform, i.e., the moment generating function, of the probability distribution for the serial interval (SI) of the disease [14, 16], denoted by *h*(*k*) and *k* is the mean SI. Since the transmission chain of 2019-nCoV remains unclear, we adopted the SI information from SARS and MERS, which share the similar pathogen as 2019-nCoV. We modelled *h*(*k*) as Gamma distributions with mean of 7.6 days and standard deviation (SD) of 3.4 days for MERS [17], and mean of 8.4 days and SD of 3.8 days for SARS [18] as well as their average, see the row heads in Table 1 for each scenario.

**Table 1.** The summary table of the estimated basic reproduction number, *R*_0_, under different scenarios. The estimated *R*_0_ is shown as in the ‘median (95%CI)’ format. The ‘reporting rate increased’ indicated the number of fold increase in the reporting rate from January 17, when WHO released the official diagnostic protocol [8], to January 20, 2020. The highlighted results are considered as the main results.

| Reporting rate increased | Estimated $R_0$ | | |
| --- | --- | --- | --- |
| | same as MERS SI<br>7.6 ± 3.4 | SI in average<br>8.0 ± 3.6 | same as SARS SI<br>8.4 ± 3.8 |
| (unchanged) | 5.31 (3.99-6.96) | 5.71 (4.24-7.54) | 6.11 (4.51-8.16) |
| 0.5-fold | 4.52 (3.49-5.76) | 4.82 (3.69-6.20) | 5.14 (3.90-6.67) |
| 1-fold | 4.01 (3.17-5.02) | 4.26 (3.34-5.38) | 4.53 (3.51-5.76) |
| 2-fold | 3.38 (2.75-4.12) | 3.58 (2.89-4.39) | 3.77 (3.02-4.67) |
| 4-fold | 2.73 (2.31-3.22) | 2.86 (2.40-3.39) | 3.00 (2.50-3.58) |
| 8-fold | 2.16 (1.90-2.45) | 2.24 (1.96-2.55) | 2.32 (2.02-2.66) |

*Note:* The ‘SI’ is serial interval. The ‘MERS’ is the middle east respiratory syndrome, and the ‘SARS’ is the severe acute respiratory syndrome.

## Results and discussion

The exponential growth fitting results are shown in Fig 1(b), (d), (f), (h), (j) and (l). The coefficient of determination, R-squared, ranges from 0.91 to 0.92 for all reporting rate changing scenarios, which implies that the early outbreak data were largely following the exponential growth. In Table 1, we estimated that the *R*_0_ ranges from 2.24 (95%CI: 1.96-2.55) to 5.71 (95%CI: 4.24-7.54) associated with 8-fold to 0-fold increase in the reporting rate. All *R*_0_ estimates are significantly larger than 1, which indicates the potential of 2019-nCoV to cause outbreaks. Since the official diagnostic protocol was released by WHO on January 17 [8], an increase in the diagnosis and reporting of 2019-nCoV infections probably occurred. Thereafter, the daily number of newly reported cases started increasing around January 17, see Fig 1, which implies that more infections were likely being diagnosed and recorded. We suggested that changing in reporting might exist, and thus it should be considered in the estimation, i.e., 8-, 4- and 2-fold changes are more likely than no change in the reporting efforts. Although six scenarios about the reporting rate were explored in this study, the real situation is difficult to determine given limited data and (almost) equivalent model fitting performance in terms of R-squared. However, with increasing reporting rate, we found the mean *R*_0_ is likely to be around 2 and 3.

Our analysis and estimation of *R*_0_ rely on the accuracy of the SI of 2019-nCoV, which remains unknown as of January 25. In this work, we employed the SIs of SARS and MERS as approximations to that of 2019-nCoV. The determination of SI requires the knowledge of the chain of disease transmission that needs sufficient number of patient samples and periods of time for follow-up [19], and thus this is unlikely to be achieved shortly. However, using SIs of SARS and MERS as approximation could provide an insight to the transmission potential of 2019-nCoV at the early stage of the outbreak. We reported that the mean *R*_0_ of 2019-nCoV is likely to be from 2.24 (8-fold) to 3.58 (2-fold), and it is largely in the range of those of SARS, i.e., 2-5 [11, 18, 20], and MERS, i.e., 2.7-3.9 [12].

We note that WHO reported the basic reproduction number for the human-to-human (direct) transmission ranged from 1.4 to 2.5 [21], which is marginally lower than ours. However, many of existing online preprints estimate the mean *R*_0_ ranging from 2 to 5 [3, 22–24], which is largely consistent with our results.

## Conclusion

We estimated the mean *R*_0_ of 2019-nCoV ranging from 2.24 (95%CI: 1.96-2.55) to 3.58 (95%CI: 2.89-4.39) if the reporting effort has been increased by a factor of between 8- and 2-fold after the diagnostic protocol released on January 17, 2020 and many medical supplies reached Wuhan.

## Declarations

### Ethics approval and consent to participate

The ethical approval or individual consent was not applicable.

### Availability of data and materials

All data and materials used in this work were publicly available.

### Consent for publication

Not applicable.

## Acknowledgements

The authors would like to acknowledge anonymous colleagues for helpful comments.

## Disclaimer

The funding agencies had no role in the design and conduct of the study; collection, management, analysis, and interpretation of the data; preparation, review, or approval of the manuscript; or decision to submit the manuscript for publication.

## Authors’ Contributions

All authors conceived the study, carried out the analysis, discussed the results, drafted the first manuscript, critically read and revised the manuscript, and gave final approval for publication.

